# Mass Amphibian Mortality during Overwintering Related To Fungal Pathogen But Not Larval Environment: An Explanation for Declines?

**DOI:** 10.1101/290213

**Authors:** Samantha L. Rumschlag, Michelle D. Boone

## Abstract

Development of infectious diseases within hosts may be shaped by environmental conditions that cause tradeoffs in energetic demands for immune responses against demands for host growth and survival. Environmental conditions may influence these tradeoffs by affecting size of hosts, or tradeoffs may change across seasons, thereby altering the impacts of diseases on hosts. In the present study, we exposed northern leopard frog (*Lithobates pipiens*) tadpoles to varying larval environments (low leaf litter, high density of conspecifics, 40 µg/L atrazine, caged fish, or control) that influenced size at metamorphosis, a measure of host quality. Subsequently, we exposed these metamorphs of to *Batrachochytrium dendrobatidis* (Bd), a fungal pathogen linked to worldwide host population declines, at metamorphosis and/or 12 weeks later, prior to overwintering. Bd exposure dramatically reduced survival during overwintering and the effect was strongest when frogs were exposed both at metamorphosis and before overwintering. Larval environments, which determined host size, did not influence effects of Bd. Stage-structured models built to assess the impacts of Bd exposure on host populations suggest that Bd exposure at metamorphosis or before overwintering would reduce annual population growth rates by an average of 19% and 41%, respectively. Our study indicates that northern leopard frog hosts suffered little effects of Bd exposures following metamorphosis and that lower host quality did not hamper a frog’s ability to respond to Bd. Instead, we provide evidence that Bd exposure can reduce survival and result in population size reductions via reduced recruitment from overwintering mortality, providing a plausible mechanism for enigmatic declines of amphibians in temperate regions.

## Introduction

Mounting evidence points to emerging infectious diseases as a major threat to public health, global economies, and wildlife populations (Binder 1999; Daszak et al. 2000; Morens et al. 2004). Thus, if our goal is to protect populations at risk of disease outbreaks, understanding factors that determine the likelihood of host population declines is key. Environmental factors can simultaneously influence hosts and pathogens; the net effect of these forces can determine the likelihood of host population declines. For instance, across disease systems, fluctuations in environmental conditions, including temperature and resource availability, result in predictable patterns of disease driven mortality of hosts (Altizer et al. 2006). For the pathogen, environmental factors can influence survival and reproductive rates, key components of pathogen virulence and persistence of the pathogen in the host population (Voyles et al. 2012). For the host, environmental factors can determine a host’s condition, ultimately affecting a host’s ability to limit damage caused by a pathogen (Pulkkinen and Ebert 2004). For instance, when conditions are physiologically stressful, little energy may be devoted to pathogen defenses compared to basal metabolic activities including host growth and survival (Wakelin 1989, Blaustein et al. 2012). Common environmental factors that induce these tradeoffs may contribute to disease development in hosts that are otherwise not impacted by certain pathogens. Environmental factors may particularly influence the susceptibility of juvenile host life stages to pathogens, because pathogen defenses including immune responses have not fully developed in juveniles compared to adults (Koop et al. 2013).

Chytridiomycosis, which some suggest is the greatest infectious disease threat on vertebrates in modern time (Murray et al. 2011), may be amplified by environmental conditions that influence host quality and pathogen virulence. Chytridiomycosis is an infectious disease of amphibians caused by the fungal pathogen, *Batrachochytrium dendrobatidis* (hereafter, Bd). It can infect hundreds of amphibian species and is present on all continents in which amphibians exist (Olson et al. 2013). Exposure to Bd can result in decreased host growth (Caseltine et al. 2016), death (Searle et al. 2011, Wise et al. 2014), and is linked with worldwide population declines of amphibians (Berger et al. 1998, Muths et al. 2003, Lips et al. 2006). Effort has been focused on understanding the disease ecology of chytridiomycosis in regions where mortality events have been sudden and widespread like Central and South America and Australia. Comparatively, little is known about the influence of Bd on amphibian populations in temperate climates in North America like the Midwest United States, which limits our understanding of the interactions that may occur between amphibian hosts and Bd and how changes in environmental conditions may lead to disease outbreaks.

Poor host condition, driven by resource availability, may be an important predictor of outcomes to Bd or other pathogen exposures. As immune responses and overwinter survival are energetically costly processes, in temperate climates larger hosts may be better suited to sustain overwintering and pathogenic exposures. Body size can predict fitness and host condition in many vertebrates (Dobson 1992, Bachman & Widemo 1999, Shine et al. 2001), including amphibians. For instance, larger body size in amphibians is associated not only with increased overwinter survival but also with earlier time to first reproduction and increased fecundity (Semlitsch & Wilbur 1988, Schmidt et al. 2012, Earl & Whiteman 2015). If size of an individual is viewed as a measure of host condition, larger individuals are likely better adapted to sustain the negative effects of pathogen exposures though a superior ability to mount energetically costly immune responses. Yet for amphibians, probability of being infected with Bd can decrease with increasing body size (Murray et al. 2013), which points to the ability of larger individuals to better defend against pathogens.

The objective of the present research is to determine the influence of larval conditions and exposure to Bd from metamorphosis through overwintering on northern leopard frog (*Lithobates pipiens*) juveniles. Northern leopard frogs are declining in parts of their range, which could be related to Bd (Carey 1993, Rorabaugh 2005, Voordouw et al. 2010). We hypothesized that 1) suboptimal conditions in the larval environment, which impact juvenile size and condition of amphibian hosts, increase the likelihood of negative effects of Bd exposure on juveniles and 2) Bd exposure and timing of exposure alter terrestrial and overwinter survival of juvenile amphibians.

## Materials and Methods

### Animal Collection and Care

Eight partial northern leopard frog (*Lithobates pipiens*) egg masses were collected on 31 March 2013 from Talawanda High School Pond (39°29’16”N, 84°43’42”W). Egg masses were held in the laboratory until 2 April 2013 when they were moved to a climate controlled room where they were held at 18.3 °C on a 14:10 h light-dark cycle until they reached free-swimming stage (Gosner stage 25 [Gosner 1960]). Tadpoles were fed ground TetraMin tropical fish flakes (Tetra Holding) ad libitum until they were transferred on 16 April 2013 (experimental day 0) to outdoor mesocosm ponds at Miami University’s Ecology Research Center (Oxford, OH, USA). To induce different size classes at metamorphosis, we manipulated larval environmental conditions with exposures to varying conditions (control, leaf litter, density of conspecifics, atrazine exposure, caged fish) within mesocosms with 5 replicates of each treatment for a total of 25 mesocosm ponds. Treatments were randomly assigned to ponds. Each mesocosm contained 1000 L water and plankton inoculates. Ponds in the low leaf litter treatment contained 300 g leaf litter from a mixed deciduous forest, while ponds associated with all other treatments contained 1 kg leaf litter. Ponds with a high density of conspecifics contained 90 northern leopard frog tadpoles, while ponds associated with all other treatments contained 30 northern leopard frog tadpoles.

Atrazine treatments consisted of exposure to 40 µg/L of atrazine (42.2% atrazine; Drexel Chemical Company); ponds assigned to all other treatments were not exposed to atrazine. To reach treatment exposure of atrazine, we dissolved 0.948 g atrazine in 1000 mL of water and added 100 mL of the solution evenly over the surface of the pond on 23 April 2013 (experimental day 7). Atrazine treatment falls within the range of expected environmental concentrations and thus represents a realistic exposure (Fairchild et al. 1998). To confirm the initial atrazine concentration, we obtained a composite water sample that represented all five mesocosms exposed to atrazine 24 hr after application and sent it to the Mississippi State Chemical Laboratory (Mississippi State, MS, USA). Analyses resulted in a measured concentration of 53 µg/L. For caged-fish treatments, on 15 April 2013 we collected 30 bluegill sunfish (15.7±1.26 cm [mean ± std. dev.]) from Acton Lake (College Corner, OH) via electroshocking. On 23 April (experimental day 7), we added two fish per mesocosm to floating cages made from plastic baskets (64.77 x 45.42 x 25.7 cm) with plastic floats that provided cover for the fish; ponds assigned to all other treatments contained floating cages and cover floats without fish as a sham control. Fish within experimental cages were fed five to ten live northern leopard frog tadpoles weekly. Additional fish were held in two holding mesocosms (1000 L) with water and opaque foam floats that provided cover. In holding mesocosms, fish were fed earthworms ad libitum. Fish from holding mesocosms were rotated into and out of floating cages in experimental ponds weekly. Northern leopard frogs from two mesocosm ponds in the caged fish treatment developed infections of parasitic copepods (possibly *Lernea cyrinacea*), so individuals metamorphosing from these ponds were excluded from the experiment. Tadpoles were reared in mesocosms through metamorphosis at which point they were transferred to the laboratory. A subsample of northern leopard frogs within each treatment was used in the terrestrial phase of the study.

During the terrestrial phase of the experiment that began after metamorphosis and lasted for 12 weeks (until experimental day 343), we housed northern leopard frogs individually within terraria that consisted of 2 L beakers containing layers of pea gravel (∼2.5 cm) and topsoil (∼4 cm) and a small water dish with a fiberglass screen attached to the tops of beakers. Treatments in the terrestrial phase of the experiment were assigned randomly to beakers. Northern leopard frogs were held at 23°C on a 14:10 h light-dark cycle in controlled environment chambers. Metamorphs were fed increasing amounts of crickets dusted in calcium powder that varied between two 0.635 cm and four 1.27 cm crickets three times per week. After 12 weeks, beginning on 11 September 2013 (experimental day 148), we initiated overwintering conditions according to James (2003). In the laboratory, we gradually decreased the amount of crickets provided to the northern leopard frogs for three feeding days until no crickets were provided.

Feeding was discontinued before temperatures were dropped to allow for gut clearance and to avoid the possibility of intestinal infection during overwintering. Beginning on 14 September 2013 (experimental day 151), we gradually acclimated the northern leopard frogs to 17°C by drawing down the temperature by 1°C every day until experimental day 156; then we held frogs at 17°C until experimental day 163. On experimental day 163 at 17°C, northern leopard frogs were exposed to Bd before overwintering (see below). To initiate hibernation after exposure to Bd, northern leopard frogs were transferred to 2 L beakers with 10 cm topsoil, a layer of leaf litter, and fiberglass screen attached to the tops of beakers and moved to an environmental chamber set to 7°C on experimental day 164. Beakers were haphazardly arranged on shelves for the remainder of the experiment. Northern leopard frogs were held until experimental day 166 at 7°C, then temperature was reduced to 6°C for 3 days before it was reduced to 5°C on a 10:14 h light-dark cycle for the remainder of the experiment (25 March 2014, experimental day 343).

Each week, we sprayed containers with dechlorinated water to maintain soil moisture during overwintering.

### Experimental Design and Bd Exposures

At metamorphosis, we assigned northern leopard frogs from different larval environments (low leaf litter, high density of conspecifics, 40 µg/L atrazine, caged fish, or control [see above]) to Bd exposure at metamorphosis (present, absent) and before overwintering (present, absent). Larval environment, Bd exposure at metamorphosis, and Bd exposure before overwintering were crossed in a factorial design resulting in a total of 20 treatments. Replication of treatments was uneven because rates of survival and metamorphosis were influenced by larval treatments; therefore, each treatment was replicated between 7 and 10 times for a total of 190 experimental units. Bd treatments were assigned randomly to individual frogs within individual treatments. The experimental unit was the individual frog.

After metamorphosis on 2 July 2013 (experimental day 77) and before overwintering on 27 September 2013 (experimental day 164), we exposed individual frogs to Bd (present or absent) for 12 h. To expose frogs to Bd, we placed individuals in ventilated plastic Petri dishes with 7 mL dechlorinated water and 1 mL of the assigned treatment solution (see below). After 12 h, individuals were returned to their assigned terrarium. We cultured Bd (isolate JEL 213 isolated from *Rana muscosa* in the Sierra Nevada Mountains [USA], obtained from J. Longcore, University of Maine, Orono, ME) on 1% tryptone agar plates using standard protocols (Longcore et al. 1999). Bd zoospores were harvested using 5 mL dechlorinated water. For Bd-absent treatments, we added dechlorinated water to 1% tryptone agar plates without Bd cultures. After 30 min, we collected the water from the plates into two solutions, one containing Bd zoospores and the other that was absent of Bd zoospores. We calculated the concentrations of Bd zoospores using a hemocytometer. The Bd-present treatment solution that frogs were exposed to after metamorphosis contained 5.2 x 10^6^ zoospores/mL. The Bd-present treatment solution that frogs were exposed to before overwintering contained 2.1 x 10^6^ zoospores/mL.

For the larval stage, we measured survival in the larval environment of all tadpoles added to the mesocosms and mass and time to metamorphosis for individuals included in the terrestrial portion of the study. In the terrestrial stage, we observed survival daily, and individual frogs were weighed weekly to measure growth. For overwintering, we measured survival and mass at the conclusion of the experiment. At the end of the experiment, frogs were euthanized using a 1% solution of MS-222 (tricaine methanesulfonate).

### Statistical Analyses

We tested for the effect of the larval environment on the proportion of northern leopard frogs surviving to metamorphosis within each pond mesocosm using analysis of variance (ANOVA). We arcsine square root transformed the survival rates. Pond means of mass at metamorphosis and time to metamorphosis of individuals used in the terrestrial portion of the experiment were log transformed and analyzed with multivariate ANOVA. In analysis of survival and mass and time to metamorphosis, the experimental unit is the mesocosm pond. For all other analyses that evaluate endpoints in the terrestrial portion of the study, the experimental unit in the statistical model is the individual; frogs were housed individually in the terrestrial portion of the experiment. We tested for the effects of larval environment, exposure to Bd at metamorphosis, and the interaction of these treatments on the survival of individuals in the terrestrial environment prior to overwintering using logistic regression. We used repeated-measures ANOVA to determine the effects of larval environment, exposure to Bd at metamorphosis, and the interaction of these treatments on mass of leopard frogs through the course of the terrestrial phase of the experiment (from one week post-Bd exposure at metamorphosis through just before overwintering) using log-transformed mass of individuals as the repeated measure. In addition, to analyze the effects of larval environment, Bd exposure at metamorphosis, and the interaction of these treatments on terrestrial growth of northern leopard frogs, we used ANOVA on change in terrestrial mass (final terrestrial mass before overwintering– mass at one week post Bd exposure at metamorphosis). We examined in the influence of larval environment, Bd exposure at metamorphosis, and Bd exposure before overwintering on overwinter survival of individuals using logistic regression. Northern leopard frogs that died prior to exposure of Bd before overwintering were excluded from analyses of overwinter survival. Non-significant interaction terms were removed from this model to conserve degrees of freedom and improve statistical power. In preliminary analyses, we included the all the two-way and three-way interactions of larval environment, Bd exposure at metamorphosis, and Bd exposure before overwintering, and in this preliminary model none of the treatments or their interactions were significant. To assess the effects of treatments on size of individuals over the overwintering portion of the experiment, we used ANOVA to test for the effects of larval environments, Bd exposure at metamorphosis, Bd exposure before overwintering, and all two way interactions of larval environments and Bd exposure at metamorphosis and before overwintering on log-transformed overwinter change in mass (mass after simulated overwintering – terrestrial mass before overwintering). We did not test the three way interaction among larval environment, Bd exposure at metamorphosis, and Bd exposure before overwintering or the two-way interaction between larval environment and Bd exposure before overwintering because low overwinter survival lead to missing cell values. Pairwise differences for larval survival and mass at and time to metamorphosis were evaluated for ANOVAs using Scheffe’s multiple comparison tests. All analyses were completed using SAS 9.2 (SAS Institute, Inc., Cary, North Carolina). ANOVAs were constructed using generalized linear models (PROC GLM) with a Gaussian distribution, and results were evaluated using Type III error with α=0.05. Logistic regressions (PROC LOGISTIC) were built with a binary distribution and a logit link function, and results were evaluated using Type III analysis of effects with α=0.05.

### Stage-Structured Population Model

To model the impacts of Bd exposure on host population growth, we built stage-based Lefkovitch (Caswell 2000) annual projection matrices using three stages representing female northern leopard frog populations with a birth pulse (Biek et al. 2002) under three conditions: no exposure to Bd, exposure of recently metamorphosed frogs to Bd, and exposure of metamorphs to Bd before overwintering. Our models consisted of three life-history stages: pre-juvenile (embryo, larva, and overwintering metamorph), juvenile, and reproductive adult. The projection matrix representing a population that has not been exposed to Bd is:

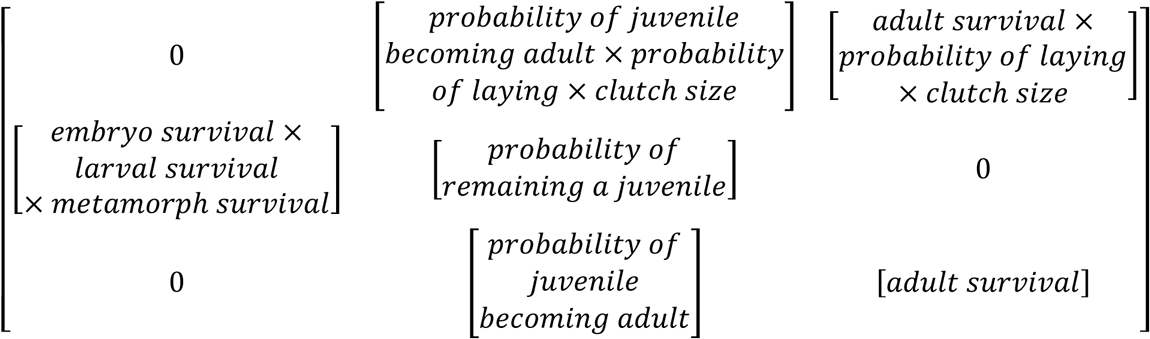

To model the effects of Bd exposure, we reduced the overwintering metamorph survival rate based on the experimental data from the current study. To represent the effects of Bd exposure at metamorphosis, we reduced overwintering metamorph survival by 42%; likewise, to represent the effects of Bd exposure prior to overwintering, we reduced overwintering metamorph survival by 77%.

Mean vital rates used in elements of the matrices were drawn from scientific literature (Table 1). When possible, means vital rates were specific to northern leopard frogs. When vital rates for northern leopard frogs were not available, we used vital rates of congeneric species. The mean embryo and larval survival rates are based off of a field survey data for wood frogs (*Rana sylvatica*) that found that survival from eggs to metamorphosis ranges from 0 to 5% (Berven 1990). The product of the mean embryo survival and the mean larval survival is the midpoint of this range, 2.5%. The mean overwintering metamorph survival and standard deviation used in the model is based on survival rates of overwintering northern leopard frog metamorphs in terrestrial enclosures where groups of 10 recently metamorphosed northern leopard frogs were overwintered in 2×2 m terrestrial enclosures with pits filled with leaf litter to allow for frogs to burrow below the frost line (Distel & Boone 2010). The mean juvenile survival rate is based on the estimates for mean juvenile survival of northern red-legged frogs (*Rana aurora*) (Biek et al. 2002).

**Table 1.** Vital rates and transition probabilities that make up the stage-structured Lefkovitch projection matrices representing females of a northern leopard frog population.

| Vital rate<sup>a</sup> | Mean | SD |
| --- | --- | --- |
| Embryo survival | 0.700<sup>(Berven 1990)</sup> | 0.049<sup>(Biek et al. 2002)</sup> |
| Larval survival | 0.036<sup>(Berven 1990)</sup> | 0.012<sup>(Biek et al. 2002)</sup> |
| Metamorph survival | 0.410<sup>(Distel and Boone 2010)</sup> | 0.050<sup>(Distel and Boone 2010)</sup> |
| Juvenile survival<sup>b</sup> | 0.335<sup>(Biek et al. 2002)</sup> | 0.093<sup>(Biek et al. 2002)</sup> |
| Juvenile to juvenile<sup>c</sup> | 0.280<sup>(Crouse et al. 1987)</sup> | 0.036<sup>(Biek et al. 2002)</sup> |
| Juvenile to adult<sup>d</sup> | 0.055<sup>(Crouse et al. 1987)</sup> | 0.056<sup>(Biek et al. 2002)</sup> |
| Adult survival | 0.178<sup>(Berven 2009)</sup> | 0.026<sup>(Berven 2009)</sup> |
| Probability of laying | 1<sup>(Biek et al. 2002)</sup> | — |
| Clutch size | 4146.50<sup>(Hupf 1977, Corn and Livo 1989, Watermolen 1995)</sup> | 856.00<sup>(Biek et al. 2002)</sup> |
| Age at sexual maturity (years) | 2 to 3<sup>(Force 1933, Vogt 1981)</sup> | — |

<sup>a</sup>Vital rates and matrix elements are for females only and are annual with the exception of embryo, larval, and metamorph survival which together represent transitions within the first year of life.
<sup>b</sup>Juvenile survival is the sum of the probability of a juvenile remaining a juvenile and the probability of a juvenile becoming an adult.
<sup>c</sup>The probability of a juvenile remaining a juvenile
<sup>d</sup>The probability of a juvenile becoming an adult

The formula used for the mean probability of a juvenile remaining a juvenile is P_1_ = ((1-p_i_^di-1^) * p_i_) / (1-p_i_^di^) where p_i_ is the annual probability of survival for a juvenile and d_i_ is the number of years spent as a juvenile (Crouse et al. 1987). We estimated female northern leopard frogs reaching sexual maturity in 2 to 3 years (Force 1933, Vogt 1981). The mean probability of a juvenile remaining a juvenile is estimated from the midpoint using the above formula with the mean juvenile survival rate and 2 years to sexual maturity and the mean juvenile survival rate and 3 years to sexual maturity. The formula used for the mean probability of a juvenile becoming an adult is P_2_ = (p_i_^di *^ (1-p_i_)) / (1-p_i_di) (Crouse et al. 1987). The mean probability of a juvenile becoming an adult is based off of the midpoint using the formula above with the juvenile survival rate with 2 years to sexual maturity and the juvenile survival rate with 3 years to sexual maturity.

The mean adult survival rate and standard deviation used in the models is based off of measurements of annual female wood frog survival in a natural population (Berven 2009). A probability of laying a clutch of 1.00 and a standard deviation of 0.000 was assumed based on a similar assumption by Biek et al. (2002) for red-legged frogs. The mean clutch size is based on the midpoint between the range of reported values of clutch sizes for northern leopard frogs from 645 to 7648 (Hupf 1977, Corn & Livo 1989, Watermolen 1995). Our model represents only female northern leopard frogs, so within the matrices, clutch size is divided by two assuming a 1:1 sex ratio.

Standard deviations for mean vital rates were taken from Biek et al. (2002) with the exception of metamorph survival and adult survival as described previously (Table 1); standard deviation for clutch size for northern leopard frogs was based on the standard deviation for boreal toads (*Anaxyrus boreas*), a species with a similar mean clutch size, and standard deviation for all other vital rates were based on a closely related species, the northern red-legged frog.

To determine the influence of reduced overwintering metamorph survival we calculated λ at stable age distribution for 2000 replicate matrices, which were generated by randomly selecting clutch sizes from a log-normal distribution and all other vital rates from β-distributions that were constructed with 2000 observations using means and standard deviations in Table 1 based on Biek et al. (2002).

Additionally, to assess the influence of vital rates on λ, the finite rate of increase of population growth, on our three annual projection matrices, we used sensitivity analysis to quantify how relatively small changes in each vital rate would affect λ when the other vital rates are held constant and elasticity analysis to quantify how proportional changes in each vital rate would affect λ when all other vital rates are held constant (De Kroon et al. 2000). Sensitivity and elasticity analyses are based on mean vital rates (Table 1). We compared sensitivity and elasticity analyses across our three projection matrices to determine if Bd exposure of juveniles influences which stage of a population is more vulnerable to small fluctuations in its vital rates. Modeling exercises were completed in R version 3.2.1. with code adapted from Stevens (2010).

## Results

Larval treatments impacted survival to metamorphosis, time to metamorphosis, and size at metamorphosis, which set the stage for testing how host condition would influence the impact of Bd on the terrestrial life stage (Table 2, Figure 1). Exposure of tadpoles to high-density and low leaf litter reduced survival to metamorphosis compared to exposure to 40 µg/L atrazine, caged fish, or the control (Figure 1a). Small to large size classes were generated by larval environmental conditions. A high density of conspecifics increased the time to metamorphosis relative to the control, caged fish, atrazine, and low leaf litter treatments (Figure 1b). Tadpoles exposed to caged fish and atrazine or control conditions reached the largest mass at metamorphosis (Figure 1b), while exposure to the low leaf litter or high density of conspecifics led to smaller sizes (Figure 1b).

**Figure 1.**
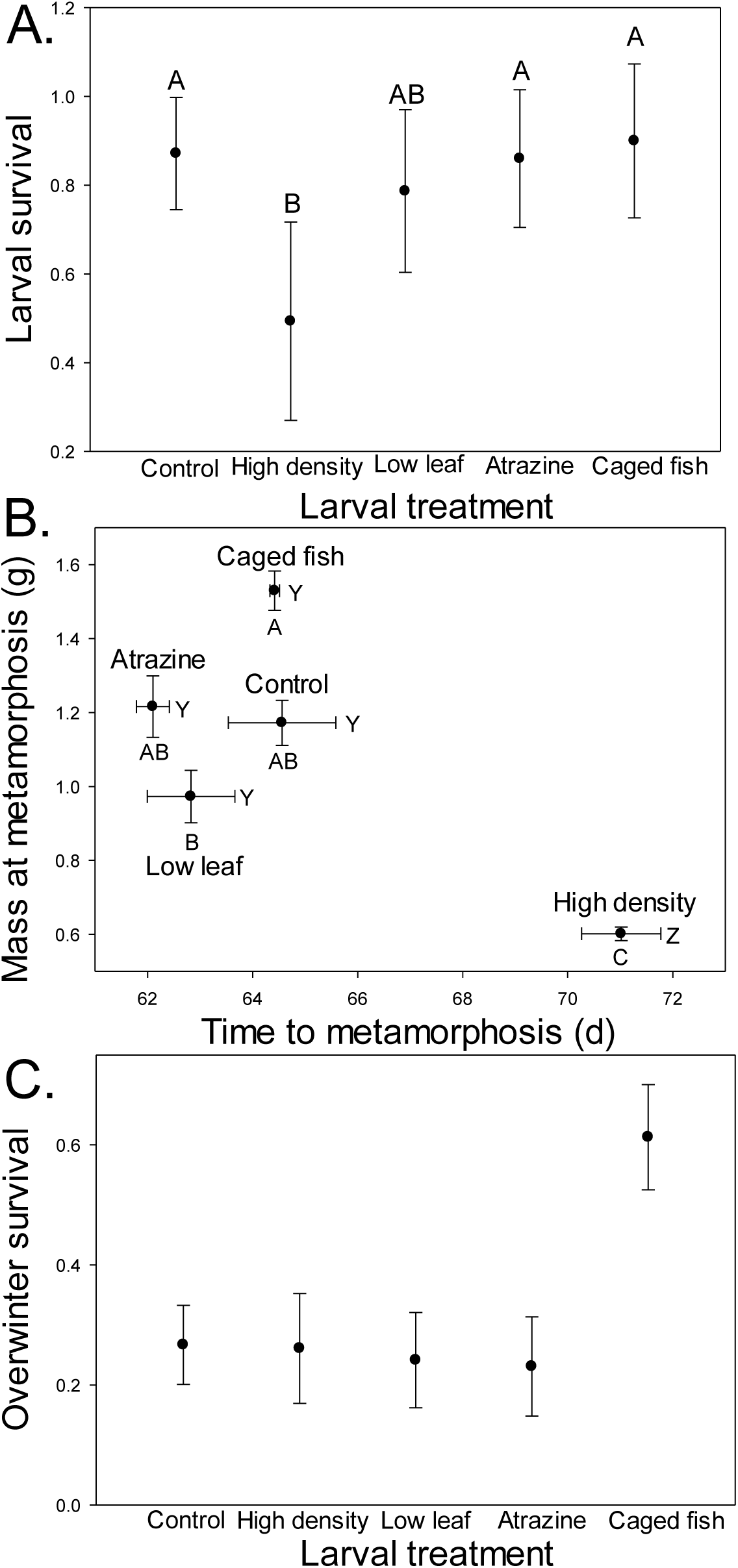
Effects of larval environment on metamorphic endpoints. Shared letters indicate no significant difference according to Scheffe’s multiple comparisons test. A.) Survival of northern leopard frog larvae exposed to varying larval environments (control, high density of conspecifics, low leaf litter, 40 µg/L atrazine, caged blue gill). Plotted values are means ± 1 binomial SE. B.) Time to metamorphosis and mass at metamorphosis for northern leopard frog larvae exposed to varying larval environments (control, high density of conspecifics, low leaf litter, 40 µg/L atrazine, caged blue gill). WXYZ corresponds with differences among treatments for time to metamorphosis. ABCD corresponds with differences among treatments for mass at metamorphosis. Plotted values are means ± 1 SE. C.) Overwinter survival of northern leopard frog metamorphs exposed to varying larval environments (control, high density of conspecifics, low leaf litter, 40 µg/L atrazine, caged blue gill). Plotted values are means ± 1 binomial SE.

**Table 2.** Summary of analyses on survival to metamorphosis and metamorphic responses of northern leopard frogs (*Lithobates pipiens*) in response to exposure to varying larval environments (control, high density of conspecifics, low leaf litter, 40 µg/L atrazine, caged blue gill).

| Response | Source of Variation | df | F | P |
| --- | --- | --- | --- | --- |
| Survival to metamorphosis | Larval environments | 4 | 7.16 | 0.0009 |
|  | Error | 20 |  |  |
| Metamorphic response | Larval environments | 8, 38 | 15.41 | <0.0001 |
| Time to metamorphosis | Larval environments | 4 | 16.99 | <0.0001 |
|  | Error | 20 |  |  |
| Mass at metamorphosis | Larval environments | 4 | 32.40 | <0.0001 |
|  | Error | 20 |  |  |

**Table 3.**
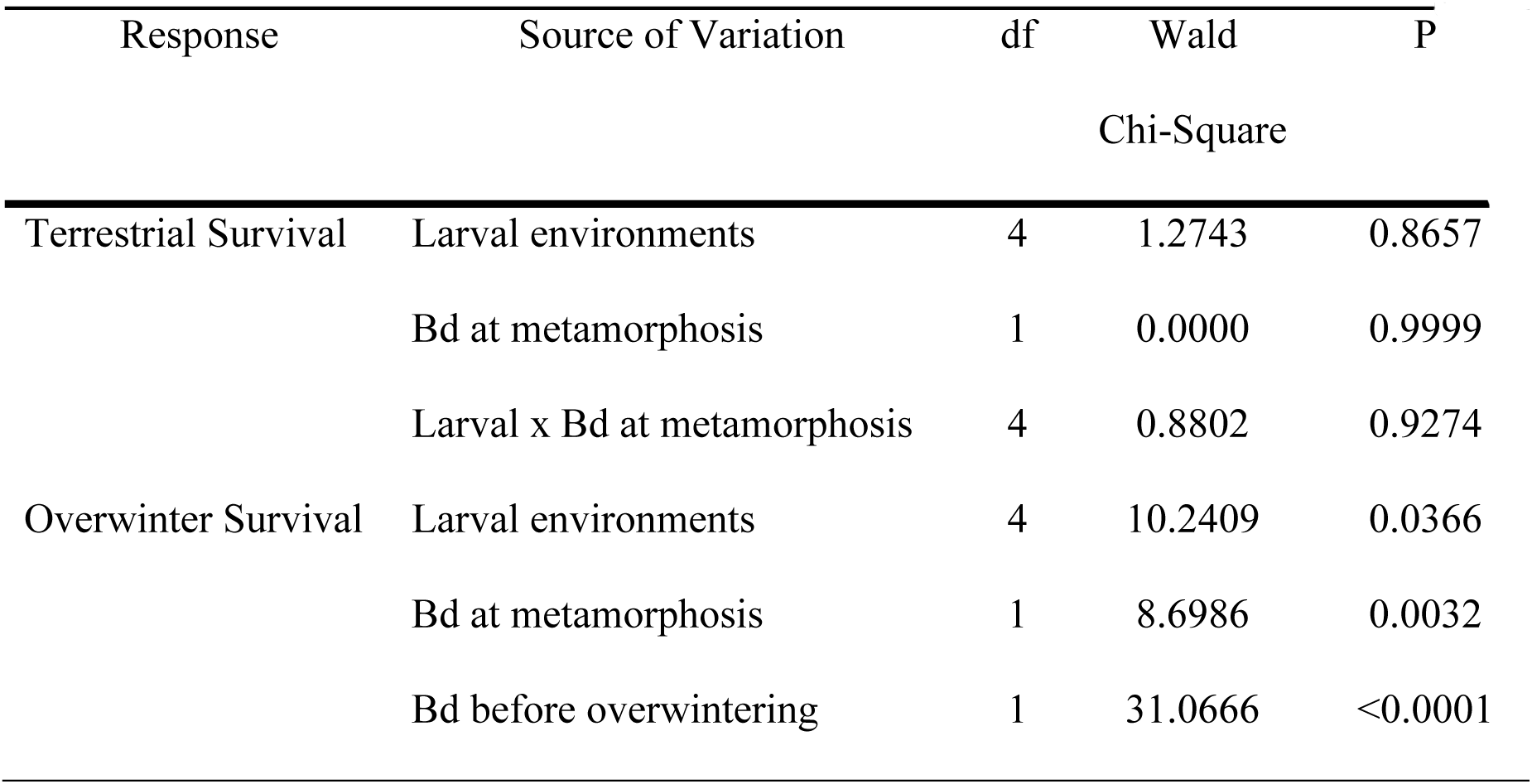
Summary of ANOVAs of northern leopard frogs (*Lithobates pipiens*) for 1) terrestrial survival in response to varying larval environments (control, high density of conspecifics, low leaf litter, 40 µg/L atrazine, caged blue gill), Bd exposures at metamorphosis (present, absent) and the interaction of these treatments and 2) survival after overwintering in response to varying larval environments (control, high density of conspecifics, low leaf litter, 40 µg/L atrazine, caged blue gill), Bd exposures at metamorphosis (present, absent), and Bd exposures just before overwintering (present, absent).

### Hypothesis I: Suboptimal Larval Conditions Increases the Likelihood of Negative Effects of Bd Exposure

Even though larval condition did have long-term effects on juveniles in that initial differences at metamorphosis were maintained throughout the terrestrial period (Figure 2), individuals exposed to different larval conditions were not differentially impacted by Bd exposure in growth or survival during the period before overwintering (i.e., no larval environment by Bd interactions, Tables 4 and 5).

**Figure 2.**
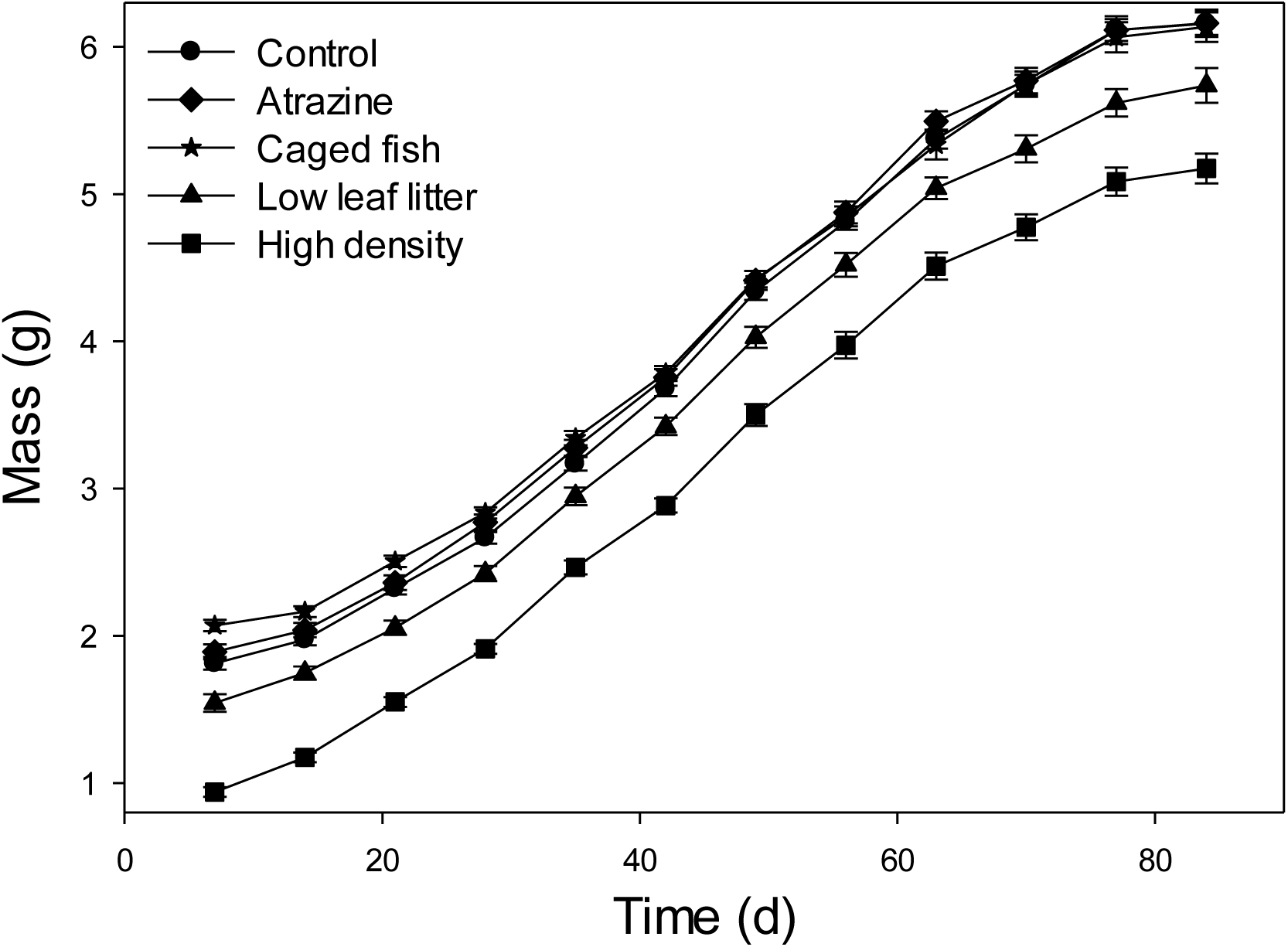
Growth of northern leopard frog metamorphs over the terrestrial portion of the experiment. Mass of northern leopard frog metamorphs over time in the terrestrial phase according to varying larval environments (control, high density of conspecifics, low leaf litter, 40 µg/L atrazine, caged blue gill). Plotted values are means ± 1 SE.

**Table 4.**
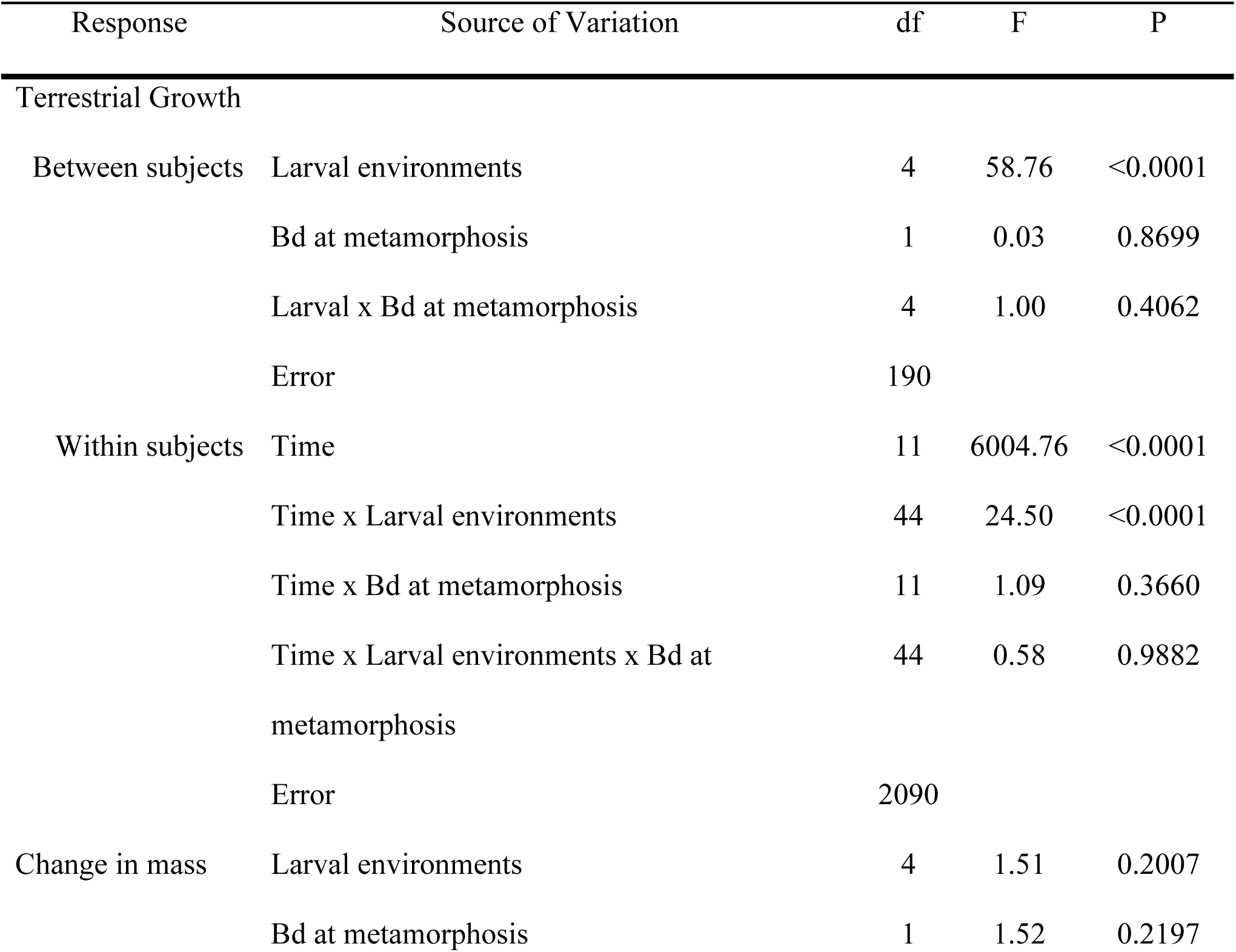
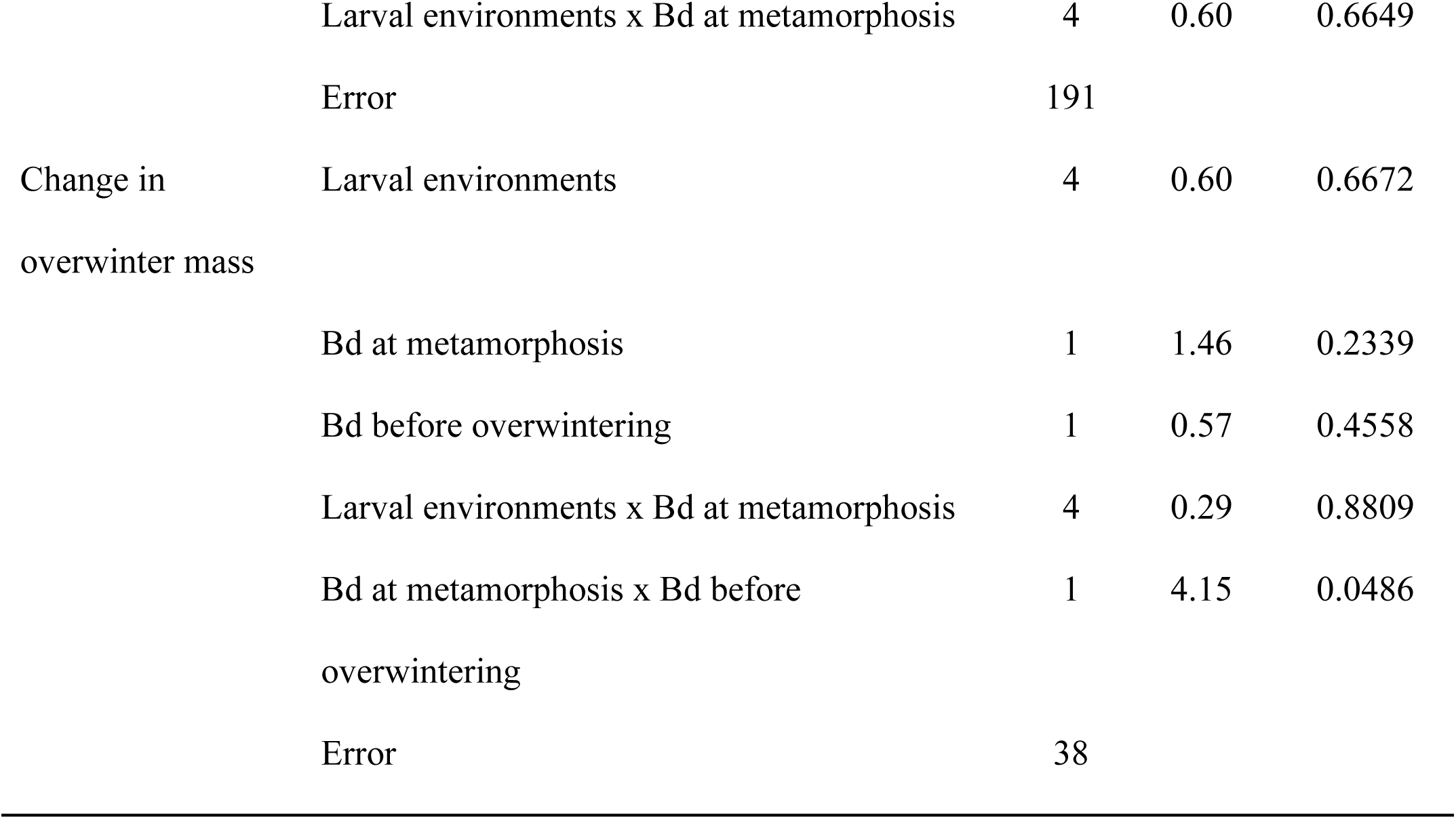
Summary of 1) repeated-measures ANOVA on juvenile mass through time and 2) ANOVA of change in mass of northern leopard frogs (*Lithobates pipiens*) over the terrestrial portion of the experiment in response to exposure to varying larval environments (control, high density of conspecifics, low leaf litter, 40 µg/L atrazine, caged blue gill), to Bd at metamorphosis (present, absent), and the interactions of these treatments. Additionally, 3) summary of ANOVA on change in mass of northern leopard frogs over the overwintering portion of the experiment in response to exposure to varying larval environments (control, high density of conspecifics, low leaf litter, 40 µg/L atrazine, caged blue gill), to Bd at metamorphosis (present, absent), to Bd before overwintering (present, absent), and the interactions of these treatments.

**Table 5.** Sensitivity and elasticity values for projection matrices representing no Bd exposure, Bd exposure of recently metamorphosed frogs, and Bd exposure of metamorphs before overwintering. The first numbers listed in a column refers to sensitivity values, and the second refers to elasticity values.

| *No Bd Exposure* |  |  |  |
| --- | --- | --- | --- |
|  | Pre-juvenile | Juvenile | Adult |
| Pre-juvenile | - | 0.004, 0.356 | 0.000, 0.056 |
| Juvenile | 52.479, 0.412 | 0.523, 0.111 | - |
| Adult | - | 1.333, 0.056 | 0.064, 0.009 |
| *Bd Exposure at Metamorphosis* |  |  |  |
| Pre-juvenile | - | 0.003, 0.325 | 0.000, 0.066 |
| Juvenile | 68.795, 0.390 | 0.531, 0.140 | - |
| Adult | - | 1.263, 0.066 | 0.079, 0.013 |
| *Bd Exposure of Metamorphs before Overwintering* |  |  |  |
| Pre-juvenile | - | 0.002, 0.263 | 0.000, 0.082 |
| Juvenile | 109.261, 0.344 | 0.549, 0.205 | - |
| Adult | - | 1.114, 0.082 | 0.107, 0.025 |

### Hypothesis II: Bd exposure and timing of exposure alters terrestrial and overwinter survival of juvenile amphibians

Timing of Bd exposure had a strong effect on the likelihood of surviving the winter (Table 3). Northern leopard frogs never exposed to Bd had a 73% probability of surviving the winter, while those exposed to Bd at metamorphosis had a 42% reduction in survival. Individuals exposed immediately prior to overwintering, however, had a 77% reduced probability of surviving the winter compared to controls (Figure 3). Likewise, larval conditions also influenced overwinter survival with individuals raised with caged fish, which were the largest at metamorphosis, having the highest survival probability (Table 3, Figure 1c). Bd exposure influenced mass after overwintering, as well (Table 4). In the absence of Bd exposure at metamorphosis, frogs lost more mass when they were exposed to Bd just before overwintering. However, this effect was reversed when frogs were exposed to Bd at metamorphosis: frogs gained more mass when exposed to Bd at metamorphosis and before overwintering compared to exposure only at metamorphosis, but low survival during overwintering resulted in low replication in some treatments (Figure 4).

**Figure 3.**
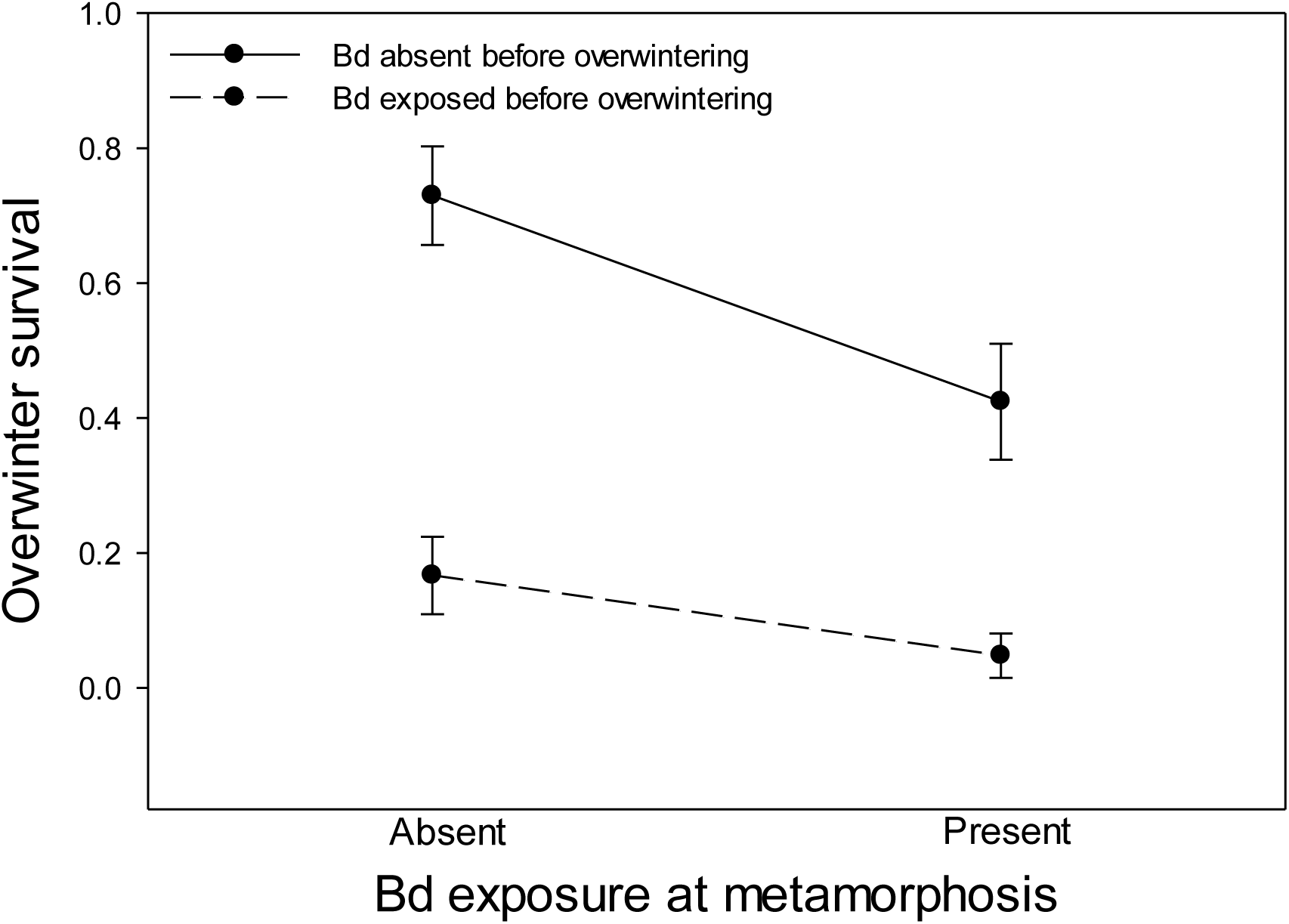
Overwinter survival of northern leopard frog metamorphs exposed to Bd at different time points, at metamorphosis (present, absent) and before overwintering (present, absent). Plotted values are means ± 1 binomial SE.

**Figure 4.**
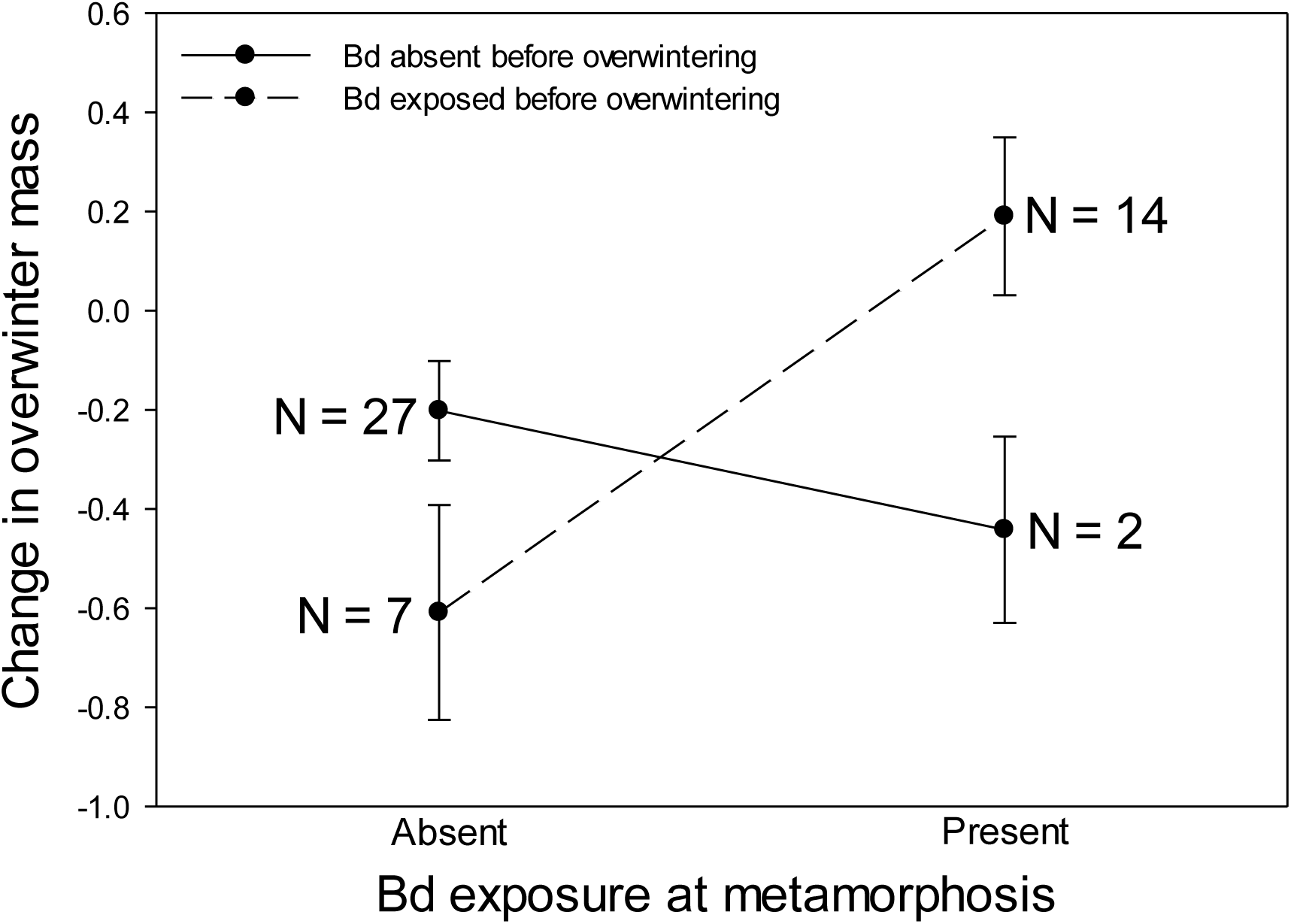
Influence of treatments on overwinter change in mass (mass after simulated overwintering – terrestrial mass before overwintering) of northern leopard frog metamorphs exposed to Bd at metamorphosis (present, absent) and before overwintering (present, absent). Plotted values are means ± 1 SE. Overwinter mortality lead to differences in number of replicates per treatment, which are shown in the figure.

### Population Model

Exposures of metamorphs to Bd (modeled as decreases in overwintering metamorph survival) had differential effects on the finite rate of increase of population growth, λ. In the model that represented a population that has not been exposed to Bd, mean λ was 1.14. When recently metamorphosed frogs were exposed to Bd, which decreased their overwinter survival and probability of transitioning to the juvenile life stage by 42%, mean λ decreased by 19% (Figure 5). When metamorphic frogs were exposed to Bd just before overwintering, which decreased their overwinter survival and probability of transitioning to the juvenile stage by 77%, mean λ decreased by 41% relative to the model with no Bd exposure (Figure 5).

**Figure 5.**
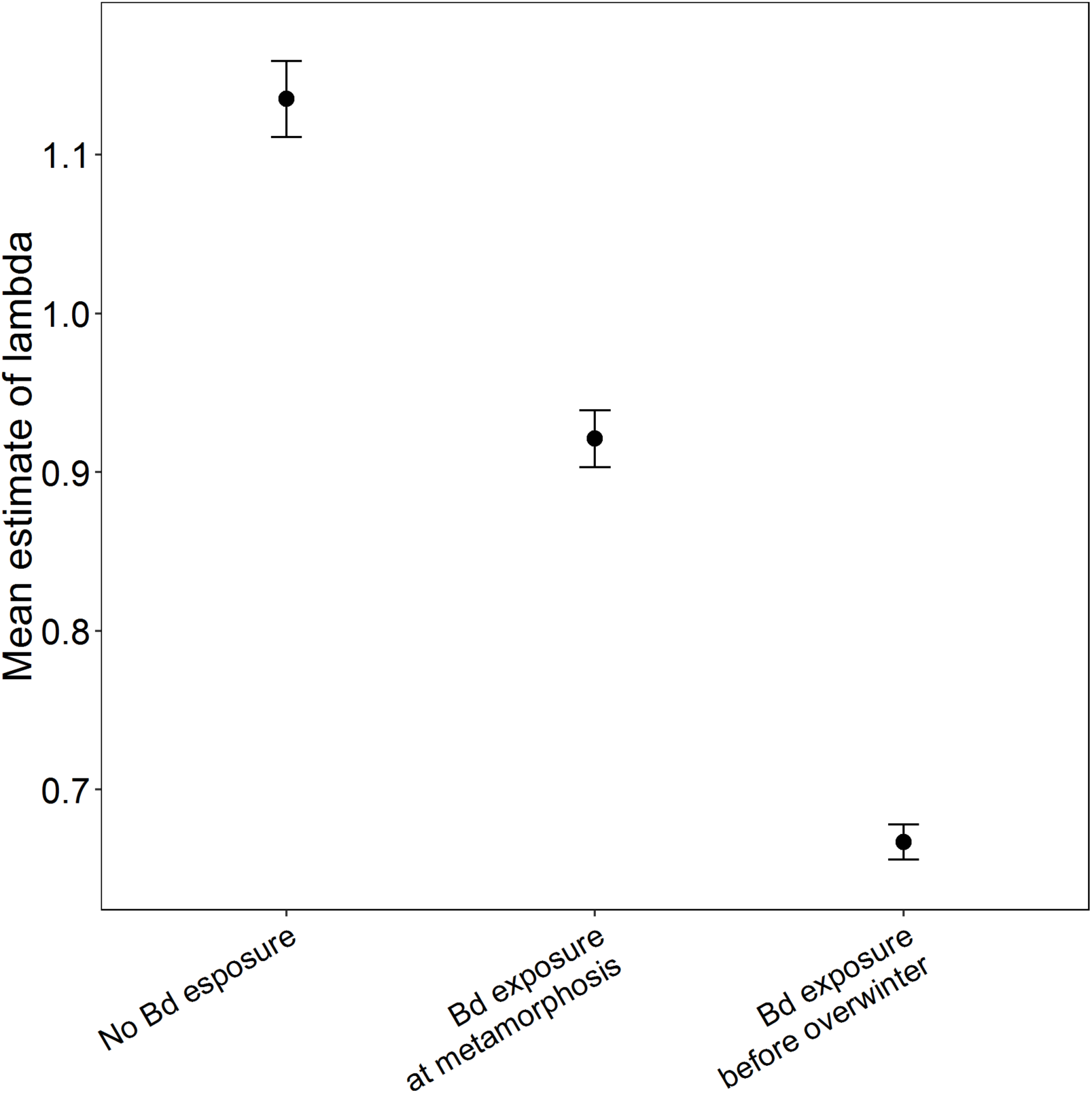
Mean λ values with 95% confidence intervals representing the impact of Bd exposures on overwinter metamorph survival to the juvenile stage

Sensitivity analyses across the three annual projection matrices, representing populations of northern leopard frogs with no exposure to Bd, exposure of recently metamorphosed frogs to Bd, and exposure of metamorphs to Bd before overwintering, showed that λ was most sensitive to changes in survival from the pre-juvenile (embryo, larva, and overwintering metamorph) to juvenile stage relative to other matrix elements (Table 4). Small changes in transition probability of pre-juveniles to juveniles would cause the biggest changes in λ. Similarly, elasticity analyses within each of these three projection matrices showed that λ was most elastic to changes in survival from the pre-juvenile to juvenile stage relative to other matrix elements (Table 6). Small proportional changes in this transition relative to all other matrix elements would have the greatest impact on λ.

## Discussion

Determining the influence of external abiotic factors on virulence of pathogens remains a largely unanswered question in disease ecology (Lively et al. 2014). Variation in climatic conditions associated with seasonality and in the quality of the environment that influences resource availability during early development represents two of the universal forces that can drive population dynamics (e.g., Fretwell 1972); yet, in temperate climates we lack an understanding of how host-pathogen interactions in wildlife populations respond to these changes over a given year. We set out to examine how larval condition of individuals influenced host vulnerability to pathogen exposures, yet despite large differences in metamorph size—an indicator of host quality and an indirect measure of fitness—we did not find individuals to be differentially affected by pathogen exposure. However, we did find that overwinter survival was disproportionally impacted by pathogen treatment. Our study is the first to demonstrate how the impact of pathogen exposure can appear to be “no effect” for a period of time (12 weeks) and then result in dramatically high overwinter mortality. Our models of population growth rates support the potential that decreases in overwinter survival could have negative impacts on at the population level. Although Bd has been found to cause local extinction of hosts (Skerratt et al. 2007), local extirpation during overwintering has not been directly observed and could offer an explanation for enigmatic declines especially for temperate species.

### Hypothesis I: Suboptimal Larval Conditions Increase the Likelihood of Negative Effects of Bd Exposure

Exposure to stressors during early life stages can have lasting impacts on health, increasing susceptibility to infectious pathogens during later life stages (Rohr et al. 2013). We predicted that exposure to suboptimal larval conditions could increase the effects of pathogen exposure on host growth (e.g., Caseltine et al. 2016) and survival (e.g., Burrow et al., in review), but we found no support for this effect. The lasting effects of larval condition on host body size demonstrate that varying larval conditions did indeed have effects on host quality. The different size classes of hosts that were generated by different larval conditions were sustained over the course of the experiment; yet, hosts exposed to larval conditions that resulted in smaller size did not have increased vulnerability to the effects of Bd exposures compared to hosts exposed to larval conditions that resulted in larger size as we predicted based on previous evidence of reduced immunity in smaller individuals (Dobson 1992, Bachman and Widemo 1999, Shine et al. 2001). Even using a range of environmental factors that could influence susceptibility to a pathogen regardless of size (e.g., the herbicide atrazine; Rohr et al. 2013), the conditions of the larval environment did not differentially alter the effect of Bd exposure.

Our results suggest that the ability of hosts to defend against pathogens is not impacted by larval condition to a level that would increase susceptibility in this system. Larval condition may not impact factors believed to reduce vulnerability to Bd, including rate of skin sloughing (Paetow et al. 2012) or production of antimicrobial skin peptides (Pask et al. 2012) in northern leopard frogs. Further, it should be noted that northern leopard frogs have not shown high susceptibility to Bd in many studies (e.g., Tennessen et al. 2009, Voordouw et al. 2010). Because northern leopard frogs’ mortality differed significantly among Bd treatments during overwintering, it suggests factors other than larval condition influence susceptibility to this pathogen.

### Hypothesis II: Bd Exposure and Timing of Exposure Alters Terrestrial and Overwinter Survival of Juvenile Amphibians

Host susceptibility to pathogens is known to vary throughout a given year (Altizer et al. 2006). Winter is a time of the year when immune system function, across classes of hosts, including humans, declines in response to pathogens because of decreases in resources and changes in weather conditions (Dowell 2001). Our results show decreased survival of all pathogen-exposed hosts during overwintering, which could suggest decreased or inhibited immune responses of hosts or increased pathogen virulence with lowered temperatures. Alternatively, longer periods of time may have been needed to see the lethal effects of Bd in northern leopard frogs.

However, there are reasons to believe overwintering could play a key role in disease development or susceptibility. In temperate regions, low temperatures during winter necessitate hibernation in amphibians, which initiates a series of energetically costly physiological processes that are associated with decreased immune responses (Maniero & Carey 1997, Carey et al. 1999, Rollins-Smith & Woodhams 2011) that could increase the impact of Bd on amphibians if they are exposed before overwintering. Northern leopard frogs can defend against Bd infections by sloughing epidermal layers, where Bd infection occurs (Paetow et al. 2012), and by producing antimicrobial skin peptides that inhibit Bd (Pask et al. 2012). These responses may be reduced or inhibited during overwintering because of reductions in temperature.

Timing of Bd exposure also appeared to dramatically alter overwinter mortality risk. For temperate hosts, the time just before overwintering may be a critical vulnerable window for exposure to pathogens because of changes in host immune defenses and pathogen physiology. For many parasites, development of infectious stages and population growth rates are linked to seasonal changes in temperature and precipitation (Altizer et al. 2006). In the present study, Bd exposure just before overwintering resulted in greater mortality and lower population growth rates compared to treatments when Bd exposure occurred shortly after metamorphosis. The cooler temperatures that coincide with the onset of winter may favor Bd growth and infection. Across climatic zones, Bd infections in amphibians commonly increase during months associated with decreases in temperatures relative to the annual mean (Ouellet et al. 2005, Woodhams & Alford 2005, Bosch et al. 2007), which may be caused by Bd’s thermal performance and ability to persist at low temperatures (4-14°C). In culture, across a range of temperatures from 13 to 28°C, initial release of zoospores, zoospore longevity, and maximum zoospore production is greatest at lower temperatures (13-15 °C; Stevenson et al. 2013). The balance between host immunity and pathogen may shift with decreases in temperature caused, at least in part, by Bd’s ability to thrive at low temperatures (Rollins-Smith et al. 2011), which may coincide with the onset of winter temperatures in the Midwest.

Further, exposure to Bd multiple times appeared to further diminish the probability of surviving overwintering. We observed that repeated exposures to Bd induced the greatest reductions in overwinter survival compared with all other treatments, suggesting that multiple exposures of Bd may not confer resistance to disease development when a secondary exposure occurs immediately prior to overwintering. McMahon et al. (2014) found that repeated exposures to Bd and subsequent clearance using heat treatments increased survival and induced stronger immune responses of Cuban treefrogs (*Osteopilus septentrionalis*), a species not known to become infected with Bd in natural populations. Although we did not measure infection following the initial exposure to Bd at metamorphosis, it is possible that northern leopard frogs carried low levels of Bd infection, which did not impact terrestrial growth or survival prior to overwintering but may have influenced their ability to develop an acquired immune response. While hosts may be able to behaviorally avoid Bd infection (McMahon et al. 2014) and even modify their thermoregulatory behavior to reduce likelihood of infection (Richards-Zawacki 2010), the effects of repeated exposure of Bd on survival may increase if hosts are not able to completely clear the infection or if exposure occurring just before overwintering which may reduce the ability of hosts to mount an immune response. Other studies consistently show amphibian hosts become re-infected with Bd after multiple pathogen exposures (Ramsey et al. 2010, Cashins et al. 2013, McMahon et al. 2014). Together these findings suggest that repeated exposures of Bd can continue to result in infection, which can have severe negative effects on host survival, especially if exposure occurs when conditions change impacting the ability of hosts to respond.

Based on the results of our study, we propose that pathogen exposures in temperate climates across host taxa can differentially alter components of host fitness (e.g., survival) according to season, leading to decreases in population growth rates. We hypothesize that population declines of hosts in temperate climates could result from disease development during overwintering, which is a pattern that has been observed in bats with white-nose syndrome (Langwig et al. 2015). Theoretical stage-structured population models developed based solely on reductions in overwinter survival of metamorphs with Bd exposure provide support that decreased population growth rates could result from disease during overwintering—a time when hosts are vulnerable to infectious pathogens but mass mortality would not be obvious. In this system, low overwinter survival caused by Bd may go unnoticed in natural amphibian populations because amphibians are difficult to track during the non-breeding season and reduced recruitment could be linked to a number of other plausible factors such as predation or variation in climate. Although Bd mass mortality events have not been documented in the Midwest United States, populations of amphibian hosts are commonly infected with Bd in places where declines are not apparent (Daszak et al. 2005, Longcore et al. 2007), and many Midwest amphibians are experiencing unexplained population declines, including northern leopard frogs. Northern leopard frogs have shown population declines throughout their range (Hecnar & M’Closkey 1996), most notably in the southwest (Rorabaugh 2005), which some have suggested is driven by Bd (Carey 1993, Voordouw et al. 2010). In parts of Ohio, where the current study takes place, northern leopard frog population extinctions have also been documented over the last 30 years (Jeff Davis, pers. comm.). Because of the absence of widespread mortality events in temperate zones, less attention has been given to understanding the disease ecology of chytridiomycosis compared to regions in which mortality events have been sudden and widespread like Central and South America and Australia. The results we present support a call for empirical testing and field observations of the effects of overwintering on changes in host social behavior, contact rates with infective stages, host births and deaths, and changes in host immune function and pathogen physiology to strengthen this hypothesis.

## Conclusions

In conclusion, our results show that suboptimal larval conditions do not alter the impacts of pathogen exposure on host growth and survival during the first year of life in this system.

Instead, our results suggest that hosts exposed to pathogens could suffer lower overwinter survival, which could have formative effects on population growth and persistence by reducing recruitment of breeding adults. Reduced resources and temperatures during winter may change host-pathogen interactions compared to tropical regions resulting in differences in disease dynamics that could shape amphibian populations in the Midwest, a region generally not believed to be influenced by Bd-related declines. Our results point to the need to evaluate the influence of overwintering on natural host-pathogen interactions in temperate climates, a topic that is understudied in the field of infectious disease. For hosts across taxa that overwinter in covert locations including amphibians, arthropods, reptiles, and small mammals, the impacts of pathogen exposures on host responses likely change with dramatic reductions in temperature and food availability but are difficult to discern from other causes of reduced spring recruitment like predation and climate effects. Examining the influence of pathogens across the life cycle of a host encompassing the range of environmental conditions experienced in the wild is critical if we are to understand how seasonality influences natural populations in this climatic region.

## Acknowledgments

We are grateful to M.H. Stevens for advising on our population model. Thanks to M. Youngquist, T. Hoskins, A. Gordon, M. Eyerman, J. Caseltine, C. Miekle, R. Wise, B. Mette, T. Leach, P. Garrett, P. Guiden, B. Hoven, and J. Fasciano for providing assistance with animal care and collection, and to M. Mahon and T. Leach for advice on developing code for population models. We appreciate C. Williamson and E. Overholt for providing materials and laboratory space for culturing Bd. Funding for this research was provided by the American Museum of Natural History and Miami University’s Department of Biology. The authors declare no conflicts of interest.

## Literature Cited

Altizer S, Dobson A, Hosseini P, Hudson P, Pascual M, Rohani P (2006) Seasonality and the dynamics of infectious diseases. Ecol Lett 9:467–484

Bachman G, Widemo F (1999) Relationships between body composition, body size and alternative reproductive tactics in a lekking sandpiper, the Ruff (*Philomachus pugnax*). Funct Ecol 13:411–416

Berger L, Speare R, Daszak P, Green DE, Cunningham AA, Goggin CL, Slocombe R, Ragan MA, Hyatt AD, McDonald KR, Hines HB, Lips KR, Marantelli G, Parkes H (1998) Chytridiomycosis causes amphibian mortality associated with population declines in the rain forests of Australia and Central America. Proc Natl Acad Sci U S A 95:9031–9036

Berven KA (1990) Factors affecting population fluctuations in larval and adult stages of the wood frog (Rana sylvatica). Ecology 71:1599–1608

Berven KA (2009) Density dependence in the terrestrial stage of wood frogs: Evidence from a 21-year population study. Copeia 2009:328–338

Biek R, Funk WC, Maxell BA, Mills LS (2002) What is missing in amphibian decline research: insights from ecological sensitivity analysis. Conserv Biol 16:728–734

Binder S (1999) Emerging infectious diseases: Public health issues for the 21st century. Science 284:1311–1313

Blaustein AR, Gervasi SS, Johnson PTJ, Hoverman JT, Belden LK, Bradley PW, Xie GY (2012) Ecophysiology meets conservation: Understanding the role of disease in amphibian population declines. Philos Trans R Soc B Biol Sci 367:1688–1707

Bosch J, Carrascal LM, Durán L, Walker S, Fisher MC (2007) Climate change and outbreaks of amphibian chytridiomycosis in a montane area of Central Spain: Is there a link? Proc Biol Sci 274:253–260

Carey, C. 1993. Hypothesis concerning the causes of the disappearance of boreal toads from the mountains of Colorado. Conserv Biol 7:355–362.

Carey C, Cohen N, Rollins-Smith L (1999) Amphibian declines: An immunological perspective. Dev Comp Immunol 23:459–472

Caseltine J, Rumschlag S, Boone MD (2016) Terrestrial growth of northern leopard frogs reared in the presence or absence of predators and exposed to the amphibian chytrid fungus at metamorphosis. J Herpetol 50:404–408

Cashins SD, Grogan LF, McFadden M, Hunter D, Harlow PS, Berger L, Skerratt LF (2013) Prior Infection Does Not Improve Survival against the Amphibian Disease Chytridiomycosis. PLoS One 8

Caswell H (2000) Matrix population models: Construction, analysis, and interpretation, 2nd edition. Sinauer Associates, Sunderland, Massachuetts

Corn PS, Livo LJ (1989) Leopard frog and wood frog reproduction in Colorado and Wyoming. Northwest Nat 70:1–9

Crouse D, Crowder L, Caswell H (1987) A stage-based population model for loggerhead sea turtles and implications for conservation. Ecology 68:1412–1423

Daszak P, Cunningham AA, Hyatt AD (2000) Emerging infectious diseases of wildlife - threats to biodiversity and human health. Science 287:443–449

Daszak P, Scott DE, Kilpatrick AM, Faggioni C, Gibbons JW, Porter D (2005) Amphibian population declines at Savannah River Site are linked to climate, not chytridiomycosis. Ecology 86:3232–3237

Distel CA, Boone MD (2010) Effects of aquatic exposure to the insecticide carbaryl are species-specific across life stages and mediated by heterospecific competitors in anurans. Funct Ecol 24:1342–1352

Dobson FS (1992) Body mass, structural size, and life-history patterns of the Columbian ground squirrel. Am Nat 140:109–125

Dowell SF (2001) Seasonal variation in host susceptibility and cycles of certain infectious diseases. Emerg Infect Dis 7:369–374

Earl JE, Whiteman HH (2015) Are commonly used fitness predictors accurate? A meta-analysis of amphibian size and age at metamorphosis. Copeia 103:297–309

Fairchild JF, Ruessler DS, Carlson AR (1998) Comparative sensitivity of five species of macrophytes and six species of algae to atrazine, metribuzin, alachlor, and metolachlor. Environ Toxicol Chem 17:1830–1834

Force ER (1933) The age of the attainment of sexual maturity of the leopard frog *Rana pipiens* (Schreber) in northern Michigan. Copeia 3:128–131

Fretwell SD (1972) Populations in a seasonal environment. Princeton University Press, Princeton, New Jersey

Gosner KL (1960) A simplified table for staging anuran embryos larvae with notes on identification. Herpetologica 16:183–190

Hecnar SJ, M’Closkey RT (1996) Regional dynamics and the status of amphibians. Ecology 77:2091–2097

Hupf TH (1977) Natural histories of two species of leopard frogs, Rana blairi and Rana pipiens, in a zone of sympatry in northeastern Nebraska. University of Nebraska

James S (2003) Methods for testing the combined effects of contamination and hibernation on terrestrial amphibians. In: Linder G, Krest S, Sparling D, Little E (eds) Multiple Stressor Effects in Relation to Declining Amphibian Populations. ASTM International, West Conshohocken, PA, p 169–183

Koop JAH, Owen JP, Knutie SA, Aguilar MA, Clayton DH (2013) Experimental demonstration of a parasite-induced immune response in wild birds: Darwin’s finches and introduced nest flies. Ecol Evol 3:2514–2523

Kroon H De, Groenendael J Van, Ehrlén J (2000) Elasticities: A review of methods and model limitations. Ecology 81:607–618

Langwig, K. E. et al. 2015. Host and pathogen ecology drive the seasonal dynamics of a fungal disease, white-nose syndrome. Proc Roy Soc London B 282:20142335.

Lips KR, Brem F, Brenes R, Reeve JD, Alford RA, Voyles J, Carey C, Livo L, Pessier AP, Collins JP (2006) Emerging infectious disease and the loss of biodiversity in a Neotropical amphibian community. Proc Natl Acad Sci U S A 103:3165–3170

Longcore JR, Longcore JE, Pessier AP, Halteman WA (2007) Chytridiomycosis widespread in anurans of northeastern United States. J Wildl Manage 71:435–444

Longcore JE, Pessier AP, Nichols DK (1999) *Batrachochytrium dendrobatidis* gen. et sp. nov., a chytrid pathogenic to amphibians. Mycologia 91:219–227

Maniero GD, Carey C (1997) Changes in selected aspects of immune function in the leopard frog, *Rana pipiens*, associated with exposure to cold. J Comp Physiol B 167:256–263

McMahon T a., Sears BF, Venesky MD, Bessler SM, Brown JM, Deutsch K, Halstead NT, Lentz G, Tenouri N, Young S, Civitello DJ, Ortega N, Fites JS, Reinert LK, Rollins-Smith LA, Raffel TR, Rohr JR (2014) Amphibians acquire resistance to live and dead fungus overcoming fungal immunosuppression. Nature 511:224–227

Morens DM, Folkers GK, Fauci AS (2004) The challenge of emerging and re-emerging infectious diseases. Nature 430:242–249

Murray KA, Retallick RWR, Puschendorf R, Skerratt LF, Rosauer D, McCallum HI, Berger L, Speare R, VanDerWal J (2011) Assessing spatial patterns of disease risk to biodiversity: Implications for the management of the amphibian pathogen, *Batrachochytrium dendrobatidis*. J Appl Ecol 48:163–173

Murray KA, Skerratt LF, Garland S, Kriticos D, McCallum H (2013) Whether the weather drives patterns of endemic amphibian chytridiomycosis: A pathogen proliferation approach. PLoS One 8: e61061

Muths E, Corn PS, Pessier AP, Green DE (2003) Evidence for disease-related amphibian decline in Colorado. Biol Conserv 110:357–365

Olson DH, Aanensen DM, Ronnenberg KL, Powell CI, Walker SF, Bielby J, Garner TWJ, Weaver G, Fisher MC (2013) Mapping the global emergence of *Batrachochytrium dendrobatidis*, the amphibian chytrid fungus. PLoS One 8:e56802

Ouellet M, Mikaelian I, Pauli BD, Rodrigue J, Green DM (2005) Historical evidence of widespread chytrid infection in North American amphibian populations. Conserv Biol 19:917–928

Paetow LJ, Daniel McLaughlin J, Cue RI, Pauli BD, Marcogliese DJ (2012) Effects of herbicides and the chytrid fungus *Batrachochytrium dendrobatidis* on the health of post-metamorphic northern leopard frogs (*Lithobates pipiens*). Ecotoxicol Environ Saf 80:372–380

Pask JD, Woodhams DC, Rollins-Smith LA (2012) The ebb and flow of antimicrobial skin peptides defends northern leopard frogs (*Rana pipiens*) against chytridiomycosis. Glob Chang Biol 18:1231–1238

Ramsey JP, Reinert LK, Harper LK, Woodhams DC, Rollins-Smith LA (2010) Immune defenses against *Batrachochytrium dendrobatidis*, a fungus linked to global amphibian declines, in the South African clawed frog, *Xenopus laevis*. Infect Immun 78:3981–3992

Richards-Zawacki CL (2010) Thermoregulatory behaviour affects prevalence of chytrid fungal infection in a wild population of Panamanian golden frogs. Proc R Soc B Biol Sci 277:519–528

Rohr JR, Raffel TR, Halstead NT, McMahon TA, Johnson SA, Boughton RK, Martin LB (2013) Early-life exposure to a herbicide has enduring effects on pathogen-induced mortality. Proc R Soc B Biol Sci 280:1–8

Rollins-Smith LA, Ramsey JP, Pask JD, Reinert LK, Woodhams DC (2011) Amphibian immune defenses against chytridiomycosis: Impacts of changing environments. Integr Comp Biol 51:552–562

Rollins-Smith LA, Woodhams DC (2011) Amphibian immunity: Staying in tune with the enviornment. In: Demas GE, Nelson RJ (eds) Eco-Immunology. Oxford University Press, Oxford, United Kingdom, p 92–143

Rorabaugh J (2005) Rana pipiens Schreber, 1782; northern leopard frog. In: Lannoo M (ed) Amphibian Declines: The Conservation Status of United States Species. University of California Press, Berkeley, CA, p 570–577

Schmidt BR, Hödl W, Schaub M (2012) From metamorphosis to maturity in complex life cycles: Equal performance of different juvenile life history pathways. Ecology 93:657–667

Searle CL, Gervasi SS, Hua J, Hammond JI, Relyea RA, Olson DH, Blaustein AR (2011) Differential host susceptibility to *Batrachochytrium dendrobatidis*, an emerging amphibian pathogen. Conserv Biol 25:965–974

Semlitsch RD, Wilbur HM (1988) Effects of pond drying time on metamorphosis and survival in the salamander Ambystoma talpoideum. Copeia 1988:978–983

Shine R, LeMaster MP, Moore IT, Olsson MM, Mason RT (2001) Bumpus in the snake den: Effects of sex, size, and body condition on mortality of red-sided garter snakes. Evolution 55:598–604

Skerratt LF, Berger L, Speare R, Cashins S, McDonald KR, Phillott AD, Hines HB, Kenyon N (2007) Spread of chytridiomycosis has caused the rapid global decline and extinction of frogs. Ecohealth 4:125–134

Stevens, M. H (2010) A Primer of Ecology with R. Second printing. Springer Science+Business Media, New York.

Stevenson LA, Alford RA, Bell SC, Roznik EA, Berger L, Pike DA (2013) Variation in thermal performance of a widespread pathogen, the amphibian chytrid fungus *Batrachochytrium dendrobatidis*. PLoS One 8: e73830.

Vogt RC (1981) Natural history of amphibian and reptiles of Wisconsin. The Milwaukee Public Museum and Friends of the Museum, Inc., Milwaukee, Wisconsin

Voordouw MJ, Adama D, Houston B, Govindarjulu P, Robinson J (2010) Prevalence of the pathogenic chytrid fungus, *Batrachochytrium dendrobatidis*, in an endangered population of northern leopard frogs, *Rana pipiens*. BMC Ecol 10:1–10

Watermolen DJ (1995) A key to the eggs of Wisconsin’s amphibians. Madison, Wisconsin

Wise RS, Rumschlag SL, Boone MD (2014) Effects of amphibian chytrid fungus exposure on American toads in the presence of an insecticide. Enviornmental Toxicol Chem 33:2541–2544

Woodhams DC, Alford RA (2005) Ecology of chytridiomycosis in rainforest stream frog assemblages of tropical Queensland. Conserv Biol 19:1449–1459

